# A *Drosophila* RNAi screen reveals conserved glioblastoma-related adhesion genes that regulate collective cell migration

**DOI:** 10.1101/2021.08.09.455704

**Authors:** Nirupama Kotian, Katie M. Troike, Kristen N. Curran, Justin D. Lathia, Jocelyn A. McDonald

## Abstract

Migrating cell collectives are key to embryonic development but also contribute to invasion and metastasis of a variety of cancers. Cell collectives can invade deep into tissues, leading to tumor progression and resistance to therapies. Collective cell invasion is also observed in the lethal brain tumor glioblastoma, which infiltrates the surrounding brain parenchyma leading to tumor growth and poor patient outcomes. *Drosophila* border cells, which migrate as a small cell cluster in the developing ovary, are a well-studied and genetically accessible model used to identify general mechanisms that control collective cell migration within native tissue environments. Most cell collectives remain cohesive through a variety of cell-cell adhesion proteins during their migration through tissues and organs. In this study, we first identified cell adhesion, cell junction, and associated regulatory genes that are expressed in human brain tumors. We performed RNAi knockdown of the *Drosophila* orthologs in border cells to evaluate if migration and/or cohesion of the cluster was impaired. From this screen, we identified eight adhesion genes that disrupted border cell collective migration upon RNAi knockdown. Bioinformatics analyses further demonstrated that subsets of the orthologous genes were elevated in the margin and invasive edge of human glioblastoma patient tumors. These data together show that conserved cell adhesion and adhesion regulatory proteins with potential roles in tumor invasion also modulate collective cell migration. This dual screening approach for adhesion genes linked to glioblastoma and border cell migration thus may reveal conserved mechanisms that drive collective tumor cell invasion.

## INTRODUCTION

While migrating cells contribute to many processes during embryonic development and adult wound healing, abnormal cell migration drives tumor cell invasion and metastasis. During development and in cancer, cells either migrate as single cells or as interconnected small to large groups of cells called collectives (Friedl and Gilmour 2009; Friedl et al. 2012; Scarpa and Mayor 2016; Te Boekhorst et al. 2016). Especially in cancer, cells can interconvert their modes of movement, transitioning from collective to single cell movement and back (Te Boekhorst and Friedl 2016). A wide variety of cancer cells, including breast, colorectal, and thyroid carcinomas, are now known to migrate and invade as collectives both *in vitro* and *in vivo* (Cheung and Ewald 2016; Wang et al. 2016; Kim et al. 2017; Ilina et al. 2018; Libanje et al. 2019; Padmanaban et al. 2019). Recent work has shown that tumor cell collectives promote tumor invasion and metastasis and may provide a mechanism for resistance to radiation (Aceto et al. 2014; Cheung et al. 2016; Haeger et al. 2019).

The *Drosophila* border cells, which migrate collectively during late oogenesis, are a simple and genetically tractable model to identify genes required for collective cell migration (Montell et al. 2012; Saadin and Starz-Gaiano 2016). The border cell cluster consists of 4-8 epithelial-derived follicle cells that surround a central pair of polar cells (Figure 1, A-C, and F). Individual border cells stay adhered together and their movement is coordinated as an entire unit during the 3-to 4-hour journey to the oocyte (Figure 1, A-C). Multiple studies have used border cells to identify conserved genes that contribute to the migration of a variety of cancer cells, including those that invade as collectives (Yoshida et al. 2004; Madsen et al. 2015; Stuelten et al. 2018; Volovetz et al. 2020).

**Figure 1.**
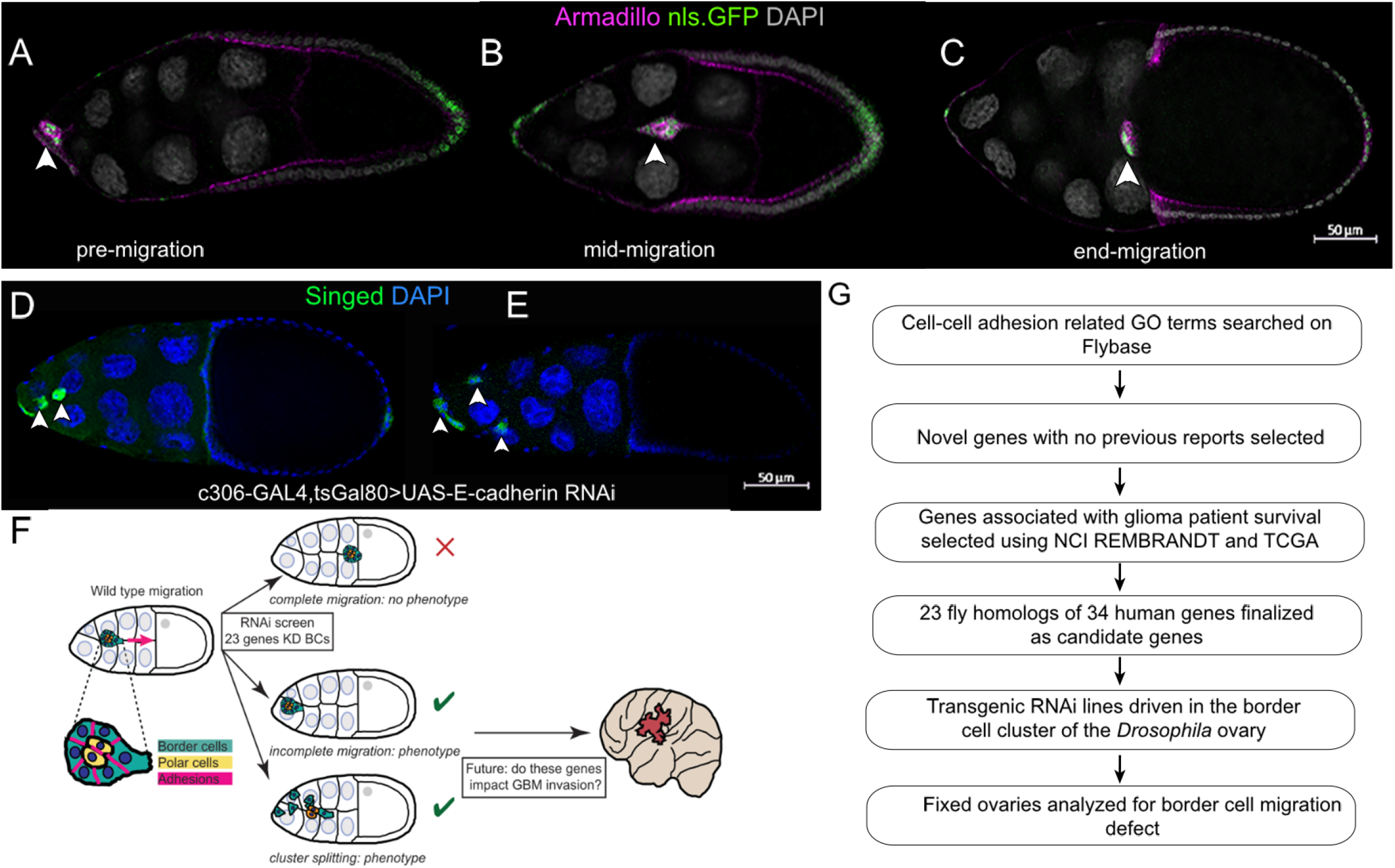
Screen to identify conserved GBM-associated adhesion genes in collective cell migration. (A-C) Migration of wild type border cells in stage 9 and 10 egg chambers. *c306*-GAL4 drives nuclear GFP (UAS-nls.GFP, green) in egg chambers labelled with Armadillo (magenta) to show cell membranes and nuclei stained for DAPI (grey). Arrowheads indicate the position of the border cell cluster within the egg chamber during migration stages: pre-migration (A), mid-migration (B), and end-migration (C). (D-E) Knockdown of *E-cadherin* by RNAi (*c306*-GAL4 tsGAL80/+; +/UAS-E-cadherin RNAi line v103962) in border cells disrupts migration and cluster cohesion at stage 10. Arrowheads indicate border cell clusters and split clusters. (F) Schematic overview of the RNAi screening approach in border cells. (G) Experimental flow chart used to identify novel GBM-associated adhesion genes through *Drosophila* and human glioma databases.

Glioblastoma (GBM) is the most common primary malignant brain tumor (Ostrom et al. 2014) and is refractory to many therapies including radiation and chemotherapy (Bao et al. 2006, Chen et al. 2012,). Given the dismal prognosis of GBM, identifying the underlying mechanisms that drive progression, including cell invasion, remains an immediate priority. While many genes are known to be dysregulated in glioma patients, it is difficult to know which ones are most relevant to disease progression, including tumor invasion. We and others recently showed that glioma cells and GBM cancer stem cells (CSCs), which can drive tumor growth, migrate collectively in some contexts (Gritsenko et al. 2017; Gritsenko and Friedl 2018; Volovetz et al. 2020). Using several patient derived GBM CSC tumor models, we found that a gene required in border cells, the small GTPase Rap1, also contributes to GBM collective cell invasion (Chang et al. 2018; Sawant et al. 2018; Volovetz et al. 2020). Because patient derived GBM CSC tumor models are less genetically accessible for screening approaches, we have turned to *Drosophila* border cells to identify conserved genes that may drive GBM collective tumor invasion but also may have a more general role in collective cell migration.

Cell-cell and cell-matrix adhesions are critical for cells to stay together and move collectively *in vivo* (Friedl and Mayor 2017; Janiszewska et al. 2020). Thus, genes that regulate cell adhesion are strong candidates to promote collective cell cohesion, migration and invasion. Here we used the border cell system to screen a subset of adhesion and adhesion-related genes that have the potential to regulate GBM tumor migration and invasion. We selected conserved adhesion genes, genes associated with cell junctions, and genes that regulate cell-cell adhesion. We further focused on those adhesion-related genes whose expression correlated with glioma patient survival but at the time of the screen did not have known functions in brain cancer. We performed an RNAi screen targeting 23 of these adhesion genes in border cells. Here, we report the identification of eight genes, *α-catenin* (*α-Cat*), *Symplekin* (*Sym*), *Lachesin* (*Lac*), *roughest* (*rst*), *dreadlocks* (*dock*), *Wnt4*, *dachsous* (*ds*), and *fat* (*ft*), whose knockdown disrupted border cell migration and/or cluster cohesion to differing degrees. We then identified three human orthologs of target genes that were enriched in the leading edge and invasive portion of GBM tumors, the α-Cat ortholog CTNNA2, the Lac ortholog NEGR1, and the Rst ortholog KIRREL3. While further work needs to be done to test these genes in GBM tumors, this study supports the use of *Drosophila* genetic approaches to provide insights into human diseases such as GBM.

## METHODS & MATERIALS

### Identification of candidate genes

FlyBase FB2014_5 version (released September 9, 2014) was queried for adhesion genes using the following Gene Ontology (GO) controlled vocabulary (CV) terms: ‘apical junction complex’, ‘focal adhesion’, ‘cell adhesion molecule binding’, ‘cell junction maintenance’, ‘cell junction assembly’, and ‘cell-cell adherens junction’. A total of 133 *Drosophila* genes were identified. Human orthologs were identified by *Drosophila* RNAi Screening Center Integrative Ortholog Prediction Tool (DIOPT) scores (Hu et al. 2011; Table 1). A PubMed search was performed for these genes along with ‘glioma’, ‘glioblastoma’, or ‘brain cancer’ to eliminate genes with a known function in or association with these cancers. This step narrowed the list to 44 genes. The NCBI REMBRANDT database was next used to identify genes that are associated with brain cancer patient survival; these results were then confirmed using The Cancer Genome Atlas (TCGA). Genes associated with better (“positive”), or worse (“negative”) patient survival were selected. These analyses resulted in 23 conserved fly genes (34 human genes) that were the final candidate genes tested in the *in vivo* border cell RNAi screen.

**Table 1.** *Drosophila* and human brain tumor-associated adhesion genes.

| *Gene name (Drosophila)* | *Human ortholog* | *DIOPT*<br>*score*<br>*out of 15* | *Best*<br>*score* | *Best*<br>*reverse*<br>*score* | *Glioma patient survival* |
| --- | --- | --- | --- | --- | --- |
| *alpha-Catenin* | CTNNA1 | 12 | No | Yes | Negative |
|  | CTNNA2 | 13 | Yes | Yes | ND |
|  | CTNNA3 | 7 | No | Yes | Positive |
| *CAP* | SORBS1 | 5 | No | Yes | Positive |
| *Caskin* | CASKIN1 | 4 | Yes | Yes | Positive |
|  | CASKIN2 | 4 | Yes | Yes | Positive |
| *CG3770* | LHFPL2 | 12 | Yes | Yes | Negative |
| *CG45049* | PERP | 10 | No | Yes | Negative |
| *Dachsous* | DCHS1 | 11 | Yes | Yes | Negative |
| *Dock* | NCK1 | 14 | Yes | Yes | Negative |
| *Fat* | FAT4 | 13 | Yes | Yes | Positive |
| *G protein alpha i* | GNAI2 | 11 | No | Yes | Negative |
|  | GNAZ | 5 | No | Yes | Positive |
| *G protein alpha o* | GNAI3 | 3 | No | No | Negative |
|  | GNAT3 | 3 | No | No | Positive |
| *Glilotactin* | NLGN2 | 5 | Yes | No | Positive |
| *Lachesin* | LSAMP | 5 | No | No | Positive |
|  | NEGR1 | 9 | Yes | Yes | Positive |
| *Liprin-alpha* | PPFIA1 | 11 | No | Yes | Negative |
| *Lowfat* | LIX1L | 13 | Yes | Yes | Positive |
|  | LIX1 | 7 | No | Yes | Positive |
| *Mesh* | SUSD2 | 13 | Yes | Yes | Negative |
| *Parvin* | PARVA | 14 | Yes | Yes | Positive |
|  | PARVB | 13 | No | Yes | Positive |
| *Roughest* | KIRREL1 | 13 | Yes | Yes | Negative |

| *Gene name (Drosophila)* | *Human ortholog* | *DIOPT score out of 15* | *Best score* | *Best reverse score* | *Glioma patient survival* |
| --- | --- | --- | --- | --- | --- |
|  | KIRREL3 | 12 | No | Yes | Negative |
|  | KIRREL2 | 11 | No | Yes | Negative |
| *schizo* | IQSEC2 | 12 | No | Yes | Positive |
| *Shroom* | SHROOM1 | 2 | No | Yes | Negative |
|  | SHROOM3 | 8 | Yes | Yes | Negative |
| *Symplekin* | SYMPK | 14 | Yes | Yes | Negative |
| *Vulcan* | DLGAP1 | 7 | Yes | Yes | Positive |
|  | DLGAP2 | 6 | No | Yes | Positive |
| *Wnt4* | WNT9A | 4 | No | Yes | Positive |
| *Wunen* | PLPP2 | 9 | No | No | Negative |

ND, not determined
Human orthologs of adhesion genes obtained from the *Drosophila* RNAi Screening Center Integrative Ortholog Prediction Tool (DIOPT). DIOPT score depicts the number of alignment tools that support an orthologous gene-pair. Best score (yes or no) indicates if the given score is the highest score. Glioma patient survival (positive or negative) is listed for each gene from The Cancer Genome Atlas (TCGA) and NCI REMBRANDT.

### Bioinformatics analyses of human genes in tumor databases

Regional gene expression data from GBM tumor tissue was obtained from the Ivy Glioblastoma Atlas Project (Ivy GAP) database (https://glioblastoma.alleninstitute.org/static/home, accessed June 20, 2021), which contains gene expression data from several anatomical features of GBM tumors in a 41 patient dataset. Analysis of gene expression based on glioma grade (grades II, III, and IV) was performed using The Cancer Genome Atlas (TCGA) data downloaded from the Gliovis data portal (http://gliovis.bioinfo.cnio.es/, accessed May 5, 2021). The GEPIA (Gene Expression Profiling Interactive Analysis; http://gepia.cancer-pku.cn/, accessed March 30, 2021) database (Tang et al. 2017) was used to compare differential expression of gene orthologs in GBM tumor tissue (n=163) and non-tumor brain tissue (n=207). Thresholds were set at a log2 fold change > 1 and a p value < 0.01.

### *Drosophila* RNAi Screen and Genetics

All genetic crosses were set up at 25°C. The tub-GAL80ts (‘tsGAL80’) transgene (McGuire et al., 2004) was included to prevent early GAL4-UAS expression and potential lethality at larval or pupal stages of development. *c306*-GAL4, tsGal80; Sco/CyO was used to drive UAS-RNAi line expression in border cells. UAS-mCherry RNAi crossed to c306-GAL4 tsGal80; Sco/CyO was used as a control. The expression pattern of *c306*-GAL4 was confirmed by crossing *c306*-GAL4, tsGal80; Sco/CyO to UAS-nls.GFP (BDSC 4776). Multiple RNAi lines for the 23 cell adhesion candidate genes and UAS-mCherry RNAi were obtained from the Vienna Drosophila RNAi Center (VDRC) or the Harvard Transgenic RNAi Project (TRiP) collection from the Bloomington Drosophila Stock Center (BDSC). All lines with stock numbers and construct IDs are listed in Table 2. Males from each UAS-RNAi line were crossed to virgin *c306*-GAL4, tsGal80 females. Three-to-five-day old F1 progeny females (*c306*-GAL4, tsGAL80/+; +/UAS-RNAi) from these crosses were fattened on wet yeast paste for ≥14 hours at 29°C prior to dissection. This allowed maximum GAL4-UAS expression and full inactivation of tsGAL80. Each RNAi line was tested one time in the primary screen, with a subset of lines tested at least three times in the secondary screen unless otherwise noted (Table 2).

**Table 2.**
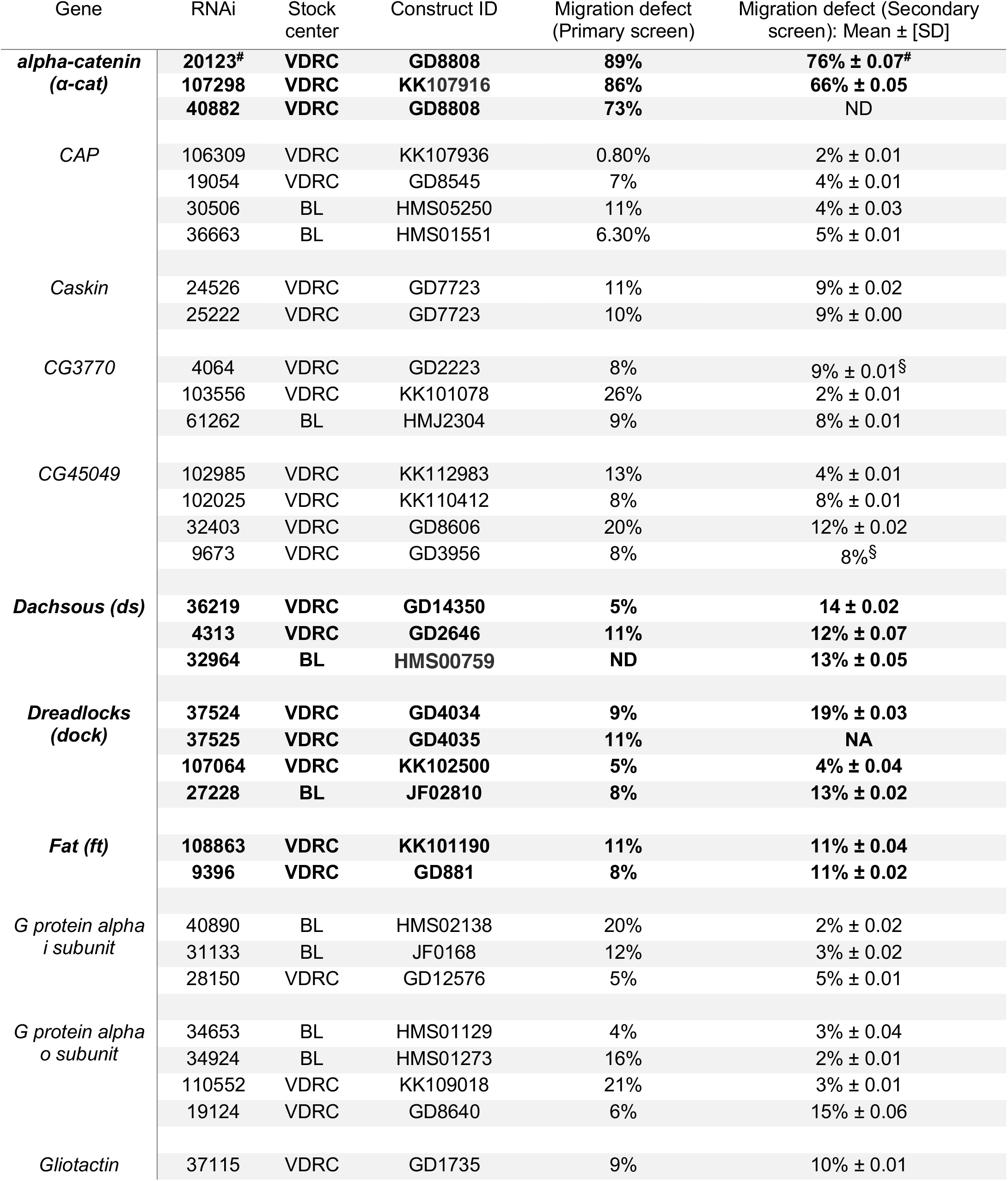
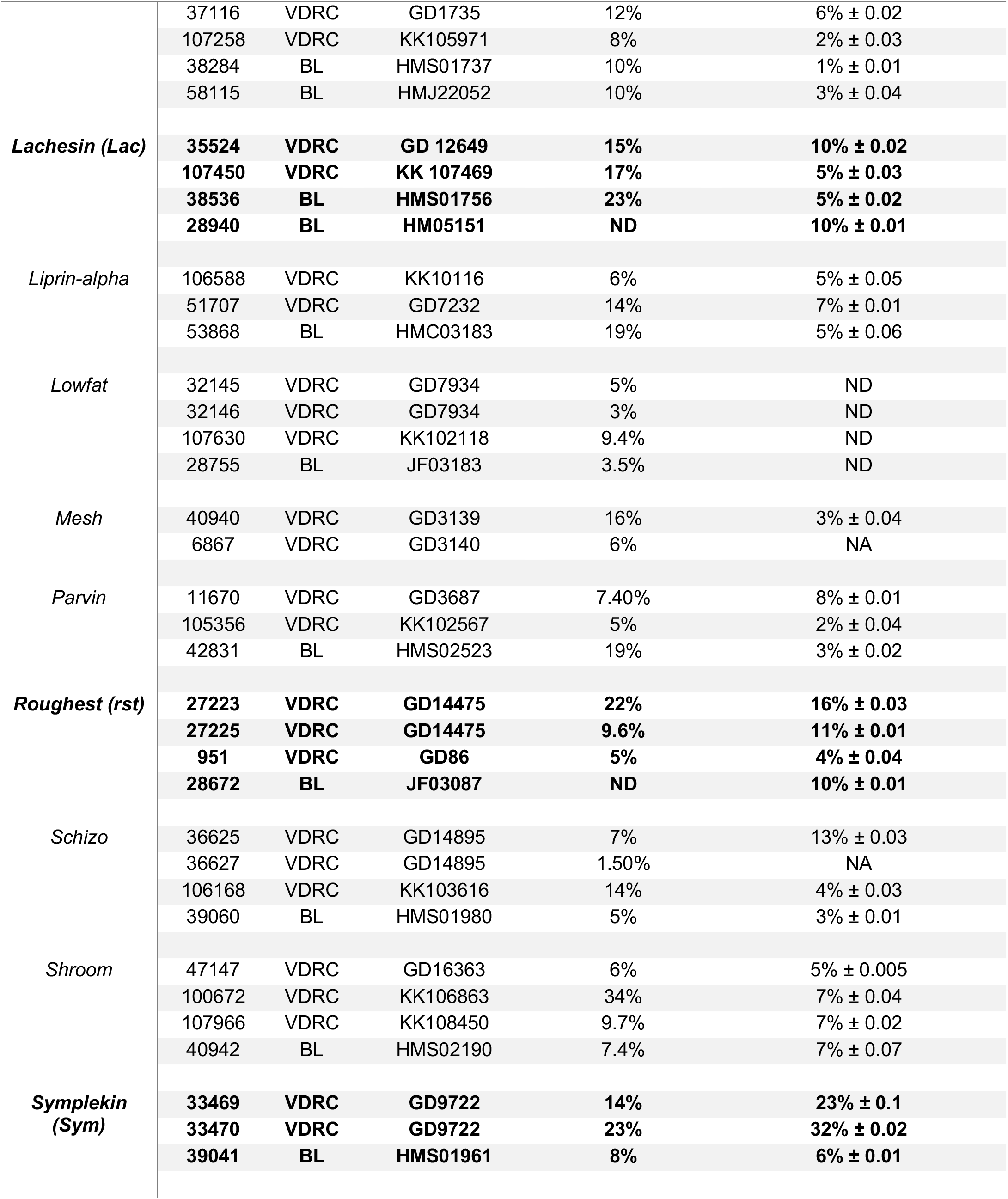
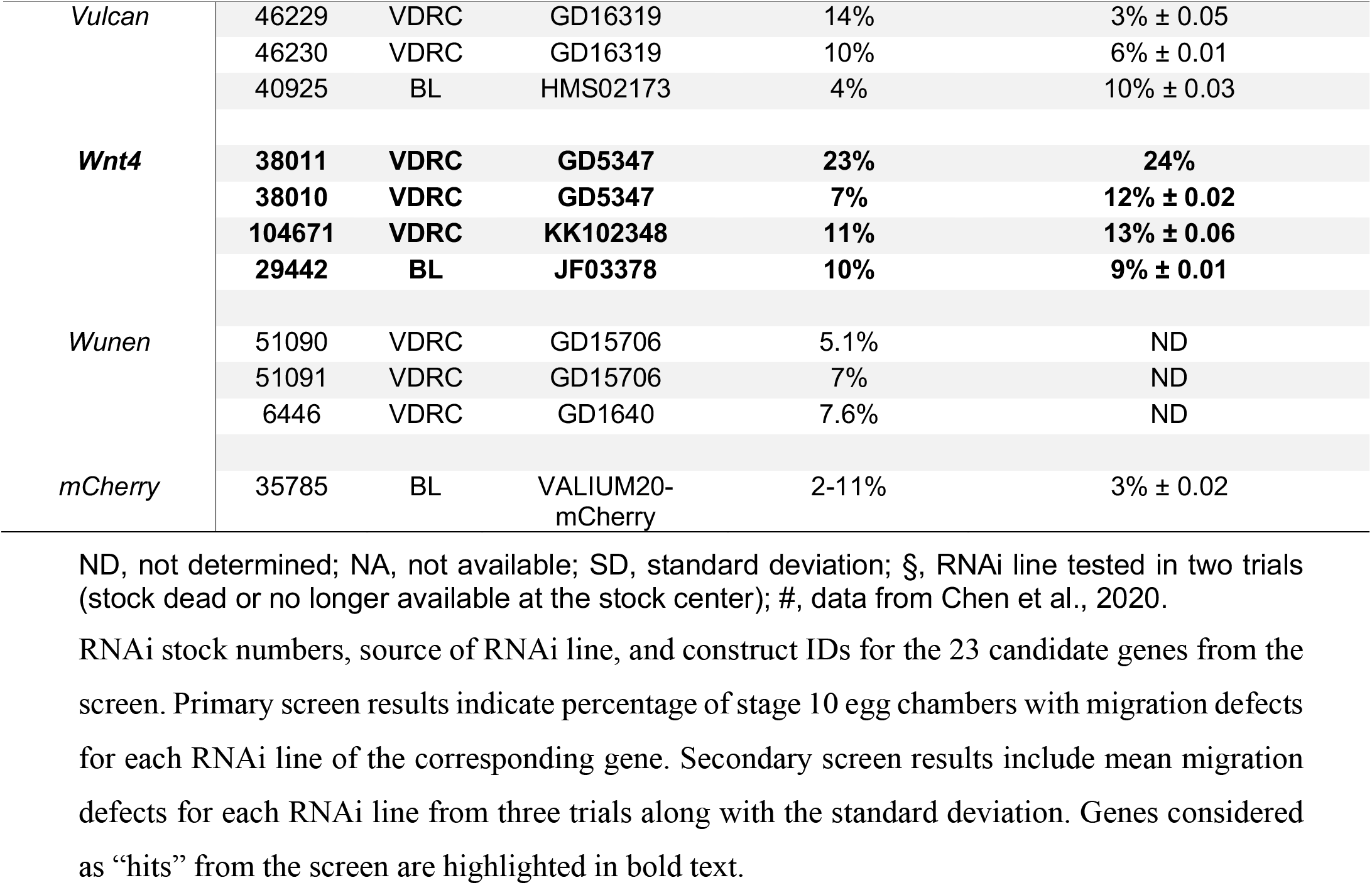
Results of the primary and secondary RNAi screens in border cells.

### Immunostaining and Imaging

Ovaries were dissected in Schneider’s *Drosophila* Medium (Thermo Fisher Scientific, Waltham, MA, USA). After dissection, ovaries were fixed in 4% formaldehyde (Polysciences, Inc., Warrington, PA, USA) in 0.1 M potassium phosphate buffer, pH 7.4 for 10 minutes. NP40 block (50 mM Tris-HCl, pH 7.4, 150 mM NaCl, 0.5% NP40, 5 mg/ml bovine serum albumin [BSA]) was used for intermediate washes and antibody dilutions. Primary antibodies were obtained from Developmental Studies Hybridoma Bank (DSHB, University of Iowa, Iowa City, IA, USA) and used at the following dilutions: rat monoclonal anti-E-Cadherin 1:10 (DCAD2), mouse monoclonal anti-Armadillo 1:100 (N27A1), and mouse monoclonal anti-Singed 1:25 (Sn7C). Anti-rat or isotype-specific anti-mouse secondary antibodies conjugated to Alexa Fluor-488 or - 568 (Thermo Fisher Scientific) were used at 1:400 dilution. 4’,6-Diamidino-2-phenylindole (DAPI, Millipore Sigma) was used at 2.5μg/ml to label nuclei. Aqua-Poly/Mount (Polysciences, Inc.) was used to mount egg chambers on slides, a coverslip was added, and the mounting media allowed to harden for three days prior to microscope imaging. The stained egg chambers were imaged either using an upright Zeiss AxioImager Z1 microscope with Apotome.2 optical sectioning or on a Zeiss LSM 880 confocal microscope (KSU College of Veterinary Medicine Confocal Core), using a 20x 0.75 numerical aperture (NA) objective. Images were processed in Zeiss ZEN 2 or FIJI software. Figures were prepared in Adobe Photoshop 2021 and line drawings were made in Adobe Illustrator 2021.

### Graphs and statistics

Graphs were prepared in GraphPad Prism 8 (GraphPad Software, San Diego, CA, USA). For the secondary screen and subsequent analyses, three trials were performed for each RNAi line (n ≥ 30 egg chambers scored in each trial). The cutoff value for a migration defect was calculated based on the background mean migration defect (3% ± 0.02) in control egg chambers (*c306*-GAL4 tsGAL80/+; +/UAS-mCherry RNAi). To determine genuine “hits” from the screen, RNAi lines with ≥10% migration defects were scored as positive hits in the primary and secondary screens. P-values were calculated using an unpaired two-tailed t test in Microsoft Excel. For GBM regional and grade-dependent gene expression analyses, differences between groups were determined using a one-way ANOVA. N’s and p-values for each trial are included in the figure legends and tables.

### Data Availability

Strains are available upon request. The authors affirm that all data necessary for confirming the conclusions of the article are present within the article, figures, and tables. Table 2 contains the complete results of the screen, including the RNAi lines tested, availability from the public stock centers (BDSC “BL” and VDRC), and detailed results from the primary and secondary screens. Supplementary Table 1 includes statistics for Figure 4 and Supplementary Figure 1. Supplementary Figure 1 shows the regional expression of the rest of the human orthologs in GBM patient tumors. Supplementary Figure 2 shows the expression of human ortholog adhesion genes in different glioma tumor grades. Supplementary Figure 3 shows a comparison of human ortholog adhesion gene expression in GBM versus non-tumor brain tissue.

## RESULTS AND DISCUSSION

### Identification of conserved brain tumor-associated adhesion genes

Cell-cell adhesion is essential for cells to stay connected during cohesive collective migration (Friedl and Mayor 2017). Reduction (or loss) of adhesion genes, such as E-cadherin (*Drosophila shotgun* [*shg*]), disrupts the integrity of the cluster and blocks the migration of the border cell cluster to the oocyte (Figure 1, D and E) (Niewiadomska et al. 1999; Sarpal et al. 2012; Desai et al. 2013; Cai et al. 2014; Chen et al. 2020, Raza et al. 2019). Many adhesion genes are conserved from flies to humans and could contribute to both border cell migration and GBM invasion (Figure 1F). To identify these conserved adhesion genes, we first performed a search of the *Drosophila* genome (FB2014_05), using Gene Ontology (GO) controlled vocabulary (CV) terms associated with cell adhesion (see Methods & Materials for details; Figure 1G). From the 133 fly genes associated with one or more of these terms, we identified likely human orthologs by analyzing their DIOPT scores (Table 1; Hu et al., 2011). Using these human orthologs, we performed a PubMed search for those genes to determine if there was an already-known association with either glioma or GBM. This allowed us to focus on genes that may have a novel association with brain tumors. The remaining 44 genes were then analyzed in the Repository of Molecular Brain Neoplasia Data (REMBRANDT), a database for transcript expression levels that are associated with brain tumor patient survival (Gusev et al., 2018). Ten genes were not found in REMBRANDT. Of the remaining 34 human genes, expression of 18 genes (13 fly genes) were associated with better (“positive”) patient survival while expression of 16 genes (13 fly genes) were associated with worse (“negative”) patient survival (Table 1). Many fly genes have multiple human orthologs. A few of these, for example *α-cat, G protein alpha i subunit*, and *G protein alpha o subunit*, have multiple human orthologs each of whose expression is associated with different predicted glioma patient outcomes (Table 1). The 23 unique fly genes were chosen for further follow-up to determine their role, if any, in border cell collective migration.

### RNAi screen in border cells identifies eight genes associated with GBM

For the primary screen, multiple RNAi lines were used to specifically target and knock down each of the 23 conserved fly adhesion genes in border cells (Table 2). We drove expression of the respective UAS-RNAi lines using *c306*-GAL4 tsGAL80, a follicle cell driver highly enriched in border cells; tsGAL80 was used to bypass early lethality (Figure 1, A-C). All border cell clusters from control (*c306*-GAL4 tsGAL80/+; +/UAS-mCherry RNAi) egg chambers completed their migration by stage 10 (Figure 2, A and B; Table 2). Twenty-one of these genes displayed a migration defect above the minimum cutoff of ≥10% with at least one RNAi line (*see* Methods & Materials).

**Figure 2.**
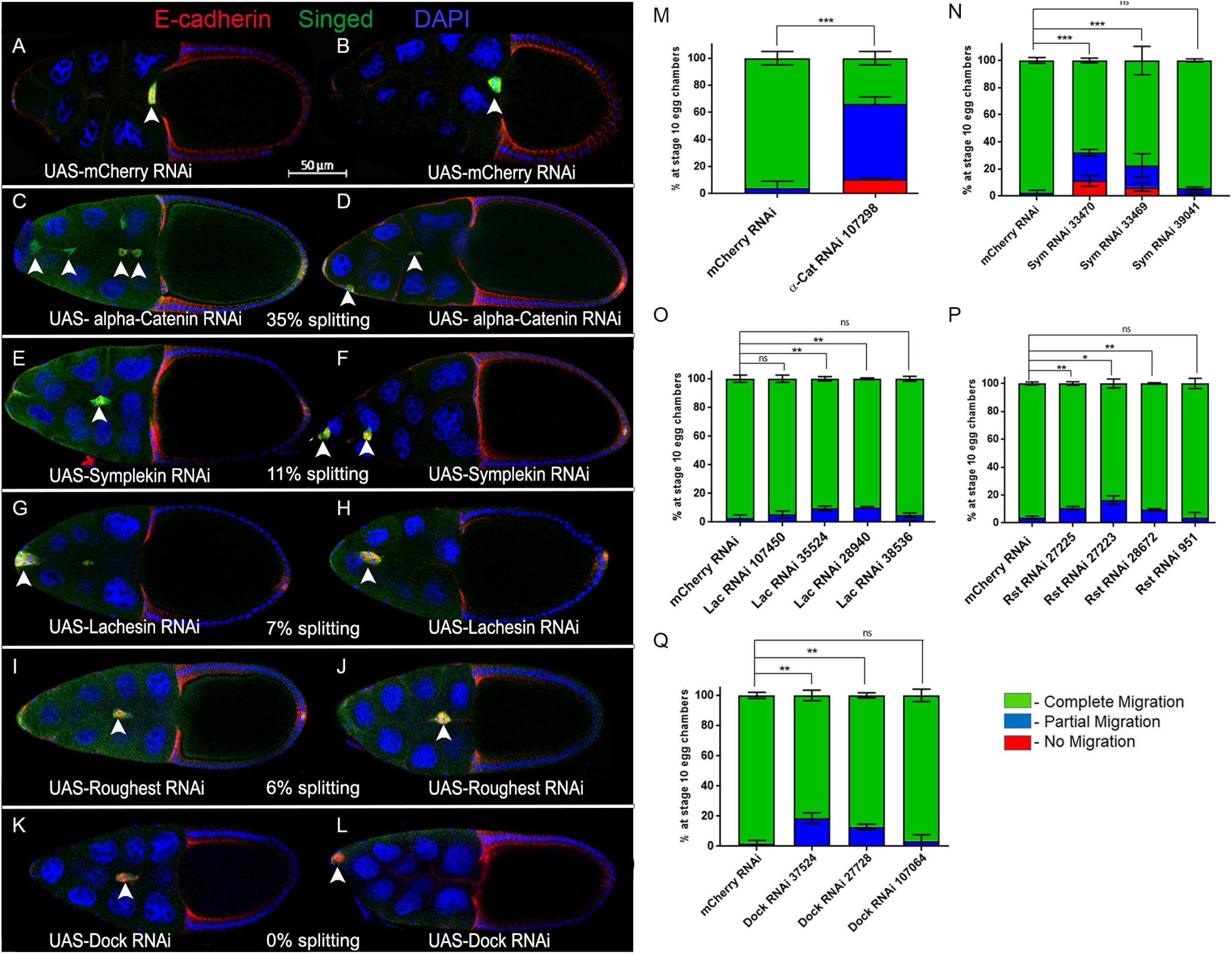
Cell adhesion and cell junction genes whose RNAi knockdown impairs border cell migration. (A-L) Stage 10 egg chambers expressing RNAi for the indicated genes (or control) in border cells labeled for E-cadherin (red), a cell membrane and adhesion marker, Singed (green), which is highly expressed in and marks border cells, and DAPI to label all cell nuclei (blue). Two images are shown to indicate the general extent of phenotypes with RNAi knockdown for each gene. White arrowheads show the position of border cell clusters; the scale bar (A, B) indicates the image magnification for all images in the figure. Anterior is to the left. (A and B) Border cells expressing the control, *mCherry* RNAi, reach the oocyte at stage 10. (C-L) RNAi knockdown of *α-Catenin/α-Cat* (C and D, line v107298), *Symplekin/Sym* (E, line v33470; F, line v33469), *Lachesin/Lac* (G and H, line BL28940), *Roughest/Rst* (I and J, line v27223) and *Dock* (K, line v37524; L, line BL27728) driven by *c306-GAL4* tsGAL80 disrupts the collective migration of border cells. The average percentage of egg chambers with border cell cluster splitting defects (% splitting) from the RNAi line with the strongest migration defect is indicated. (M-Q) Quantification of the extent of border cell migration (no migration, red; partial migration, blue; complete migration, green) in stage 10 egg chambers expressing the indicated RNAi lines for *α-Cat* (M), *Sym* (N), *Lac* (O), *Rst* (P) and *Dock* (Q) along with the matched control *mCherry* RNAi. Error bars represent SEM for three trials, n ≥ 30 egg chambers in each trial. *p<0.05; **p<0.005; ***p<0.001, unpaired two-tailed t test.

To further determine which of these genes were genuine hits, we retested the RNAi lines in a secondary screen. Each RNAi line was crossed to *c306*-GAL4 tsGAL80 three times and scored for the ability of border cells to complete their migration to the oocyte. For three genes (*ds, Lac, rst*), additional RNAi lines were obtained and tested. We specifically analyzed if RNAi border cells failed to initiate migration (“no migration”), stopped along the migration pathway but did not reach the oocyte (“partial migration”), reached the oocyte (“complete migration”), or if clusters had defective cohesion and split into multiple parts (“% splitting”). Control border cells completed their migration to the oocyte by stage 10 (Figure 2, A and B; Figure 3, A and B; Table 2). We found that knockdown of eight genes, *α-Cat, Sym, Lac, rst, dock, Wnt4, ds*, and *ft*, consistently disrupted border cell migration with at least two RNAi lines, providing more confidence that these genes are required for collective cell migration (Figures 2 and 3; Table 2). Border cell migration defects upon knockdown of these genes ranged from 10 to 76% depending on the gene and the RNAi line; some RNAi lines for these genes had less than 10% migration defects. Below we report and discuss the results for these eight genes in more detail.

**Figure 3.**
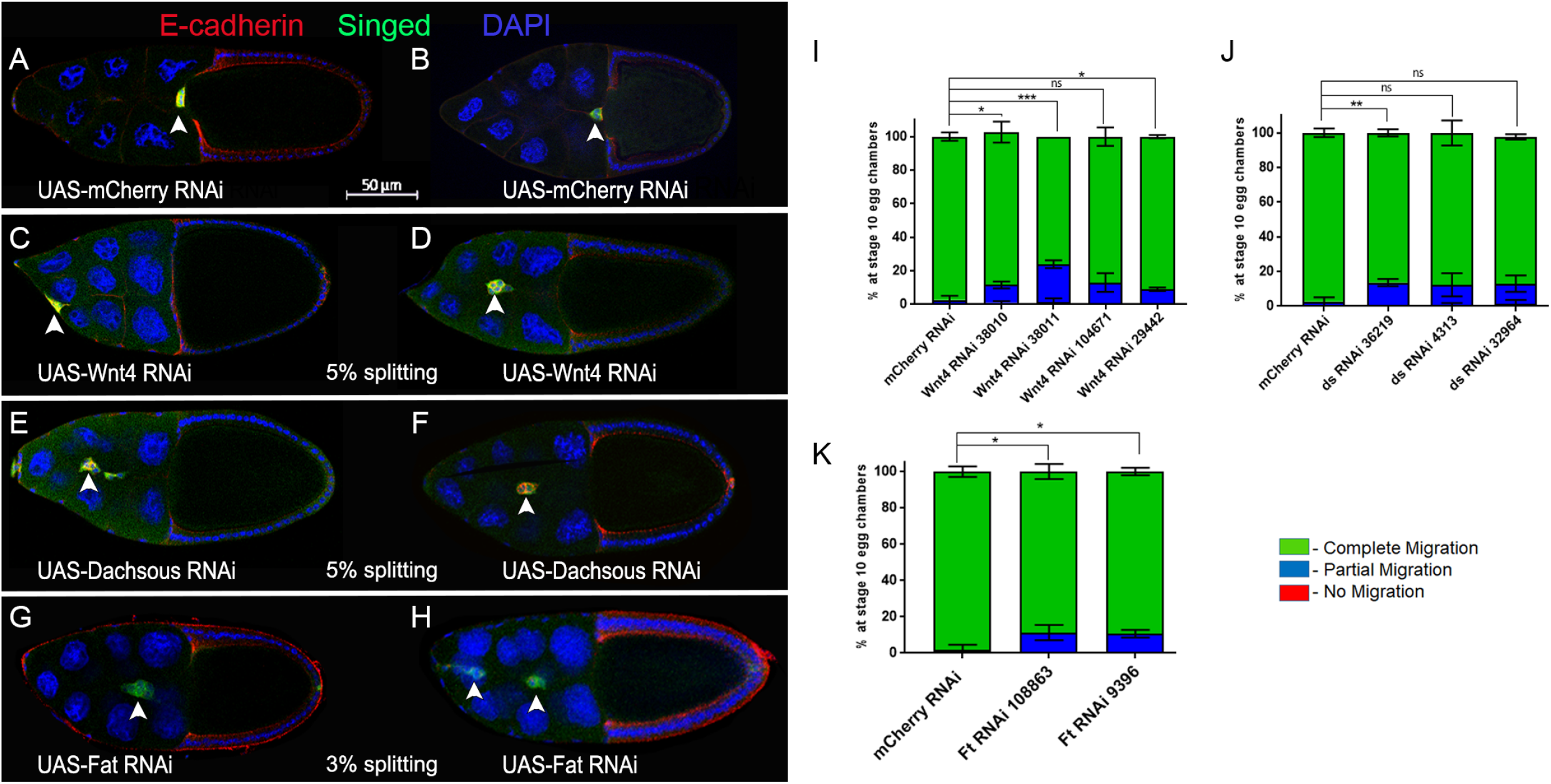
Atypical cadherins and planar cell polarity genes whose RNAi knockdown impairs border cell migration. (A-H) Stage 10 egg chambers expressing RNAi for the indicated genes (or control) in border cells labeled for E-cadherin (red), a cell membrane and adhesion marker, Singed (green), which is highly expressed in border cells, and DAPI to label all cell nuclei (blue). Two images are shown to indicate the general extent of phenotypes with RNAi knockdown for each gene. White arrowheads show the position of border cell clusters; the scale bar (A, B) indicates the image magnification for all images in the figure. Anterior is to the left. (A and B) Border cells expressing the control, *mCherry* RNAi, reach the oocyte at stage 10. (C-H) RNAi knockdown of *Wnt4* (C and D, line v38011), *Dachsous/ds* (E, line 32964; F, line v4313) and *Fat/ft* (G and H, line BL28940) driven by *c306*-GAL4 tsGAL80 disrupts the collective migration of border cells. The average percentage of egg chambers with border cell cluster splitting defects from the RNAi line with the strongest migration defect is indicated. (I-K) Quantification of border cell migration (no migration, red; partial migration, blue; complete migration, green) in stage 10 egg chambers expressing the indicated RNAi lines for *Wnt4* (I), *ds* (J), and *ft* (K) along with the matched control *mCherry* RNAi. Error bars represent SEM for three trials, n ≥ 30 egg chambers in each trial. *p<0.05; **p<0.005; ***p<0.001, unpaired two-tailed t test.

#### Adherens junction genes

α-Cat (human CTNNA1, CTNNA2, CTNNA3) is a critical component of the cadherin-catenin complex that regulates adherens junctions by linking E-cadherin and β-catenin to the F-actin cytoskeleton (Maiden and Hardin 2011). E-cadherin is required for adhesion of border cells to the nurse cell substrate, which provides traction for border cells to keep moving forward and thus facilitates forward movement while maintaining tension-based directional motility (Niewiadomska et al. 1999; Cai et al. 2014). α-Cat was the strongest candidate from our primary screen (Table 2), and we recently described the phenotypes for *α-Cat* knockdown in detail (Chen et al. 2020). *α-Cat* was knocked down using two independent RNAi lines, which reduced α-Cat protein levels in border cells (Chen et al. 2020). *α-Cat* RNAi strongly disrupted migration, with 66-76% border cells failing to complete their migration (Figure 2, C, D and M; Table 2). Border cell clusters deficient for *α-Cat* also had significant cohesion defects, with the cluster splitting into two or more parts in 35% of egg chambers (Figure 2, C and D). Thus, *Drosophila α-Cat* is required for both successful border cell migration and for proper cohesion of cells within the cluster (this study; Sarpal et al. 2012; Desai et al. 2013; Chen et al. 2020). The role for α-Cat in cluster cohesion and migration closely resembles that of β-Cat *(Drosophila* Armadillo) and E-cadherin, thus it is likely that α-Cat functions in the classical cadherin-catenin complex in border cells (Niewiadomska et al. 1999; Sarpal et al. 2012; Desai et al. 2013; Cai et al. 2014; Chen et al. 2020).

#### Other junctional genes

Four genes, *Sym, Lac, rst*, and *dock*, encode proteins that localize to various types of cell junctions and/or are known to regulate cell adhesions. Sym (human SYMPK) is a scaffolding protein, which along with other polyadenylation factors, forms a complex that mediates processing of polyadenylated and histone mRNAs but also functions at tight junctions (Keon et al. 1996; McCrea et al. 2009; Sullivan et al. 2009). During *Drosophila* oogenesis, Sym is required for histone pre-mRNA processing in the histone locus body during endoreplication of the follicular epithelium (Tatomer et al. 2014). Later in oogenesis, Sym protein localizes to the tricellular junctions of follicle cells. Here, Sym may facilitate cytoplasmic mRNA polyadenylation and thus translation of mRNAs required to regulate and/or maintain adhesion at cell junctions (Tatomer et al, 2014). Border cells expressing *Sym* RNAi had significant migration defects along with splitting of the cluster (Figure 2, E and F, N; Table 2). The two strongest *Sym* RNAi lines (VDRC 33469 and 33470), which target the same region of the *Sym* gene, caused significant migration defects, with 5-10% of border cells failing to start migration and an additional 18-22% failing to reach the oocyte. *Sym* RNAi border cell clusters had cohesion defects, with 11% of clusters visibly splitting apart. A third independent RNAi line (BL 39041) did not impair migration (Figure 2N). Based on our observed phenotypes and the known roles for Sym, we speculate that *Sym* may maintain cell-cell contacts between border cells during collective migration, possibly through regulation of as-yet-unknown targets by mRNA polyadenylation at cell-cell junctions.

Lac (human LSAMP and NEGR1) is a membrane-localized protein with three extracellular immunoglobulin-like (Ig-like) domains that can mediate cell-cell adhesion (Finegan and Bergstralh 2020). Lac localizes to both immature and mature basolateral septate junctions and is required for tracheal morphogenesis in *Drosophila* (Llimargas et al. 2004). Knockdown of *Lac* by four RNAi lines, which together target two non-overlapping regions of the *Lac* gene, mildly disrupted migration and cluster cohesion (Figure 2, G and H, O; Table 2). Two *Lac* RNAi lines (VDRC 35524 and BL 28940) disrupted migration in 11% of egg chambers, whereas two RNAi lines (VDRC 107450 and BL 38536) had fewer migration defects and were not significantly different from control (Figure 2O; Table 2). While the phenotypes caused by *Lac* RNAi knockdown are mild, *Lac* is likely to be a true regulator of border cell migration. Recent work by Alhadyian et al. found that four additional septate junction proteins, Macroglobulin complement-related (Mcr), Contactin, Neurexin-IV and Coracle, localize to border cells and are required for both border cell cluster migration and cohesion (Alhadyian et al. 2021). Because border cells do not have mature septate junctions (which form the tight occluding junctions), septate junction proteins may instead regulate cluster polarity and/or adhesion during migration (Alhadyian et al. 2021). Further work will be needed to determine if the mild phenotypes observed with *Lac* RNAi are due to partial knockdown or to redundancy with other septate junction genes.

Rst (human KIRREL1, KIRREL2, KIRREL3) is a member of the Irre Cell Recognition Module (IRM) family of transmembrane proteins. In particular, Rst encodes an immunoglobulin superfamily cell adhesion molecule (IgCAM) with five Ig-like domains (Finegan and Bergstralh 2020). IRM proteins, including Rst, control the adhesion and patterning of various tissues including the developing ommatidia in the *Drosophila* eye (Bao and Cagan 2005; Johnson et al. 2011; Finegan and Bergstralh 2020). Border cells expressing *rst* RNAi showed consistent though mild migration defects with three RNAi lines (VDRC 27223, VDRC 27225, and BL 28672), which in total target two non-overlapping regions of the *rst* gene. Migration defects ranged from 10-16% (Figure 2, I and J, P; Table 2). Cluster cohesion was mildly affected (6% of clusters split apart; Figure 2I). A fourth RNAi line did not disrupt migration or cohesion compared to control (Figure 2P; VDRC 951). Interestingly, Rst is required for progression through *Drosophila* adult oogenesis, including development of the germline (Valer et al. 2018; Ben-Zvi and Volk 2019). Rst is also expressed in follicle cells prior to the stages that border cells develop from the follicle cell epithelium (Valer et al. 2018), further supporting a later role in border cell migration.

Dock (human NCK1) is an SH2/SH3 domain-containing adaptor protein involved in receptor tyrosine kinase signaling, actin regulation, cell adhesion, and other processes (Buday et al. 2002; Chaki and Rivera 2013). In *Drosophila*, Dock regulates axon guidance, myoblast fusion during embryonic development, and ring canal morphogenesis in the ovarian germline-derived nurse cells (Garrity et al. 1996; Rao and Zipursky 1998; Kaipa et al. 2013; Stark et al. 2021). Knockdown of *dock* in border cells, using two independent RNAi lines that target non-overlapping regions of the *dock* gene (VDRC 37524 and BL 27228), resulted in migration defects but did not disrupt cohesion of border cells (Figure 2, K, L, and Q; Table 2). Specifically, *dock* RNAi disrupted migration in 13-19% of stage 10 egg chambers (Figure 2Q; Table 2). One RNAi line (VDRC 107064) did not impair border cell migration but showed mild splitting (6%), whereas another line (VDRC 37525) from the primary screen was no longer available so could not be confirmed in the secondary screen (Figure 2Q; Table 2). Dock is required for myoblast fusion during muscle formation by regulating cell adhesion and F-actin (Kaipa et al. 2013). In this context, Dock colocalizes with and/or binds to several cell adhesion proteins from the IgCAM superfamily including Rst, one of the genes identified in this screen (see above). Additionally, Dock genetically and biochemically interacts with the Ste20-like serine-threonine kinase Misshapen (Msn) to control motility of photoreceptor growth cones in the developing eye (Ruan et al. 1999). Notably, Msn is required for border cell migration, where it is required for the formation of polarized protrusions and coordinated actomyosin contractility of the cluster (Plutoni et al. 2019). Thus, it will be of interest in the future to determine if Dock, Rst, and Msn interact to control border cell migration.

#### Atypical cadherins and planar cell polarity genes

Three genes, *Wnt4, ds*, and *ft* encode proteins with annotated roles in both planar cell polarity and cell-cell adhesion (FlyBase; Figure 3; Table 2). Wnt4 (human WNT9A) is a conserved secreted protein of the Wnt family, which regulates cell adhesion through recruitment of focal adhesion complexes during the migration of epithelial cells in the pupal ovary (Cohen et al. 2002). We tested four RNAi lines for *Wnt4*, which in total target two independent regions of the gene. Migration defects for the four tested *Wnt4* RNAi lines ranged from 9 to 23% (Figure 3, C, D, I; Table 2). These data suggest a role for Wnt4 in regulating border cell movement. Previous studies suggested that Wnt4 participates in establishing planar polarity within the developing eye and wing (Lim et al. 2005; Wu et al. 2013). Indeed, several core planar cell polarity genes including *frizzled* and *dishevelled* regulate border cell migration (Bastock and Strutt 2007). However, recent studies that used multiple gene knockouts now indicate that the Wnt family of proteins, including Wnt4, are not required for *Drosophila* planar cell polarity (Ewen-Campen et al. 2020; Yu et al. 2020). Thus, we favor a role for Wnt4 in the movement and adhesion of border cells, similar to what was found during earlier stages of *Drosophila* ovarian development (Cohen et al. 2002).

Ds (human DCHS1) and Ft (human FAT4) encode large protocadherin proteins, each of which has multiple extracellular cadherin repeats (27 for Ds and 34 for Ft) (Fulford and McNeill 2020). Heterophilic binding between Ds and Ft via their extracellular domains is essential for cell-cell communication, particularly in the regulation of tissue growth through Hippo signaling and planar polarization of various tissues (Matakatsu and Blair 2004; Bosveld et al. 2016; Blair and McNeill 2018; Fulford and McNeill 2020). Knockdown of *ds* with any of three independent RNAi lines (VDRC 36219, VDRC 4313, and BL 32964) mildly disrupted migration, ranging from 12-14% of border cells failing to reach the oocyte (Figure 3, E, F, J; Table 2). *ds* RNAi border cell clusters only displayed mild cohesion defects, with 5% of clusters splitting apart (Figure 3E). Two independent RNAi lines that target *ft* (VDRC 108863 and VDRC 9396) also showed consistent though mild migration defects (11-13%), with only a few clusters (3%) splitting apart (Figure 3, G, H, K; Table 2). Interestingly, *ds* is required for the collective directional migration of *Drosophila* larval epidermal cells (LECs) during morphogenesis of the pupal abdominal epithelium (Bischoff 2012; Arata et al. 2017). An imbalance in Ds protein levels between LECs during collective migration is detected by Ft at cell junctions leading to the formation of lamellipodia at the posterior side of the LECs (Arata et al. 2017). Further experiments will be needed to determine if Ft and Ds similarly coordinate protrusions in border cells or regulate some other aspect of border cell collective migration.

### Analysis of regional expression of border cell screen hits in GBM tumors

Based on the results of the functional *Drosophila* screen, we next sought to link individual genes to invasion in human GBM patient tumors. We first assessed the Ivy GAP database that provides regional RNA expression across anatomically defined regions of tumors ranging from the tumor core to the infiltrating edge (see Methods & Materials). Using this database, we found that NEGR1 and KIRREL3 were specifically enriched in anatomical regions with elevated invasion potential, namely the leading edge (LE) and infiltrating tumor (IT), compared to all other assessed anatomical regions (Figure 4A; Supplementary Table 1). These regions included cellular tumor (CT), perinecrotic zone (PNZ), pseudopalisading cells around necrosis (PAN), hyperplastic blood vessels (HBV), and microvascular proliferation (MP). Additionally, CTNNA2 had significant expression in the LE and IT regions though was expressed in other regions of the tumor (Supplementary Figure 1; Supplementary Table 1). However, we also observed some *Drosophila* screen hits that did not demonstrate regional heterogeneity in terms of expression, such as SYMPK and CTNNA1 (Figure 4B; Supplementary Table 1). Other genes had a mixture of expression profiles across human GBM anatomical regions (CTNNA3, DCHS1, FAT4, KIRREL1, KIRREL2, NCK1; Supplementary Figure 1; Supplementary Table 1). WNT9A was not found in the Ivy GAP database. It is worth noting that this initial validation approach takes advantage of regional differences within the same GBM tumor. Therefore, such GBM anatomical expression surveys may be a better surrogate of cellular invasion than expression in GBM compared to lower-grade or non-neoplastic neural tissue; these latter analyses rely on gene expression in tissue obtained mainly from the core of the tumor and may miss areas of the tumor that undergo active invasion (Supplementary Figures 2 and 3). Nonetheless, we observed a variety of human adhesion ortholog gene-dependent increases or decreases in GBM tumors compared to lower-grade or non-neoplastic neural tissue (Supplementary Figures 2 and 3). Together, these assessments provide a first step in validating novel, conserved molecular mechanisms of GBM invasion for future therapeutic development. Invasive GBM is thought to be driven by CSCs, which can migrate and invade as single cells, finger-like collectives, or as a mixture of migration modes (Cheng et al. 2011; Volovetz et al., 2020). Human Rap1a, originally identified in a *Drosophila* screen of collective border cell migration, influences CSC-mediated GBM cell invasion (Aranjuez et al., 2012; Volovetz et al. 2020). Interestingly, knocking down *Sym* and *α-Cat* in the border cells caused the most severe migration and cluster cohesion defects. While the respective human orthologs SYMPK, CTNNA1, and CTNNA2 did not show regional tumor heterogeneity, they are each expressed in GBM tumors and/or are generally elevated in different grades of glioma including GBM (Grade IV; Supplementary Figure 2).

**Figure 4.**
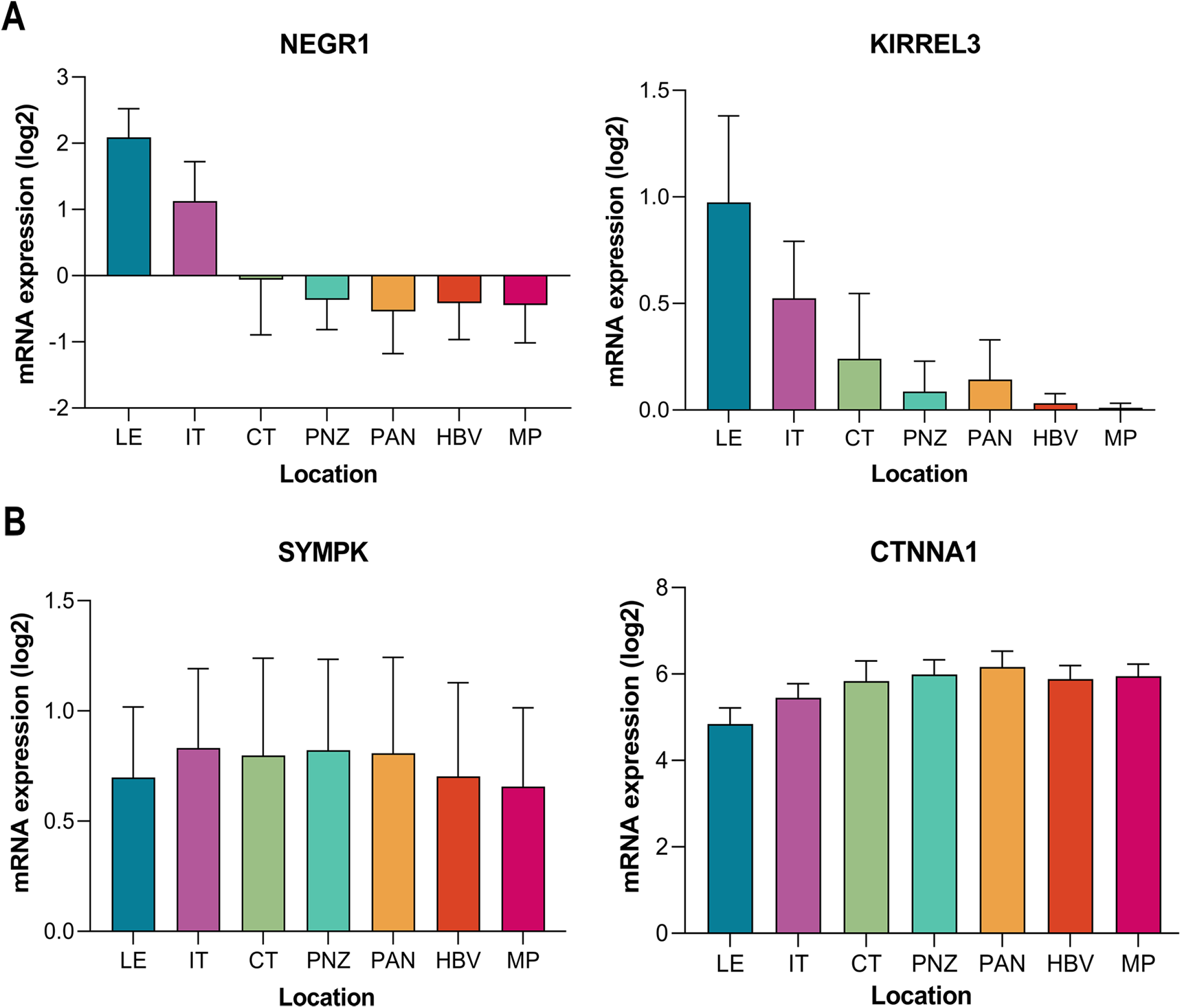
Regional expression of representative human ortholog adhesion genes in GBM patient tumors. (A) Expression of human orthologs of adhesion genes neuronal growth regulator 1 (NEGR1) and kirre like nephrin family adhesion molecule 3 (KIRREL3) is significantly enriched in the leading edge (LE) and infiltrating tumor (IT) compared to other tumor regions, including the cellular tumor (CT), perinecrotic zone (PNZ), pseudopalisading cells around necrosis (PAN), hyperplastic blood vessels (HBV), and microvascular proliferation (MP). (B) In contrast, expression of human orthologs symplekin (SYMPK) and catenin alpha 1 (CTNNA1) demonstrated little to no significant change when comparing different regions of tumor. Data from the Ivy GAP are shown as mean expression +/- SD across GBM tumor regions. Statistics are shown in Supplementary Table 1: *p<0.05; **p<0.01; ***p<0.001, one way ANOVA with Tukey HSD.

## CONCLUSION

GBM, the most common primary malignant brain tumor in adults, is also one of the most lethal (Ostrom et al. 2014; Ostrom et al. 2018). These tumors are highly invasive and possess a self-renewing CSC population. CSCs are highly invasive and can migrate either individually or collectively (Cheng et al. 2011, Volovetz et al. 2020). Here we used a human GBM-informed approach to identify conserved regulators of adhesion during collective cell migration and invasion, particularly focused on testing genes in the border cell model. We identified eight adhesion-related *Drosophila* genes (orthologs of 13 human genes) associated with glioma patient survival. Of the eight adhesion-related *Drosophila* genes found to be essential for collective cell migration, two human orthologs, NEGR1 and KIRREL3 showed significant regional enrichment in the leading edge and infiltrating tumor of human GBM tumors, areas associated with enhanced cell invasion. CTNNA2 was expressed in these invasive regions, though was also expressed at high levels in other regions of the tumor. Knockdown of these eight genes disrupted border cell migration to varying degrees, with two genes *α-cat* and *Sym* significantly disrupting both cohesion of the cluster and successful cell migration. These eight *Drosophila* genes thus represent a starting point to further investigate the specific mechanisms by which these genes regulate normal collective cell migration. Whether the human orthologs function through an adhesion-dependent or -independent manner in GBM tumors needs to be determined with follow up experiments, using both mammalian and non-mammalian models of GBM (Shahzad et al. 2021). Overall, the strategy used in this study has the potential to identify new genes and conserved mechanisms that drive collective cell migration of normal cells and those in invasive cancers such as GBM.

## ACKNOWLEDGMENTS

We thank the Vienna *Drosophila* Resource Center, Harvard Transgenic RNAi Project, and Bloomington *Drosophila* Stock Center for providing flies, and the Developmental Studies Hybridoma Bank at the University of Iowa for providing antibodies used in this study. We thank the Kansas State University College of Veterinary Medicine Confocal Core for use of the Zeiss LSM880 confocal. Thank you to C. Luke Messer, Emily Burghardt and Dr. Yujun Chen. for helpful comments on the manuscript.

---

**Supplementary Figure 1.** Regional expression of human ortholog adhesion genes in GBM patient tumors for additional human genes. See Figure 4 legend for abbreviations of the tumor regions. Data from the Ivy GAP are shown as mean expression +/- SD across GBM tumor regions. Statistics are shown in Supplementary Table 1: *p<0.05; **p<0.01; ***p<0.001, one way ANOVA with Tukey HSD.

**Supplementary Figure 2.** Expression of human ortholog adhesion genes across glioma tumor grade. Box plots of mRNA expression obtained from the TCGA database in grade II (n=226), grade III (n=244), and grade IV (n=150) patient gliomas. *p<0.05; **p<0.01; ***p<0.001, one way ANOVA with Tukey HSD.

**Supplementary Figure 3.** Expression of human ortholog adhesion genes in GBM compared to non-tumor brain tissue. Box plots of mRNA expression obtained from the GEPIA database in nontumor (n=207) and GBM tumor (n=163). *p<0.01.

**Supplementary Table 1.** Statistics for the Ivy GAP regional gene expression for all human adhesion gene orthologs. Graphed data are shown in Figure 4 and Supplementary Figure 1.

